# Nanopore sequencing reveals the DNA replication timing programme of the malaria parasite *Plasmodium falciparum*

**DOI:** 10.64898/2026.10.12.758579

**Authors:** Xiangru Huang, Francisco Berkemeier, Rosie Berners-Lee, Katy Sherborne, Sarah E. Chapman, Catherine J. Merrick, Michael A. Boemo

## Abstract

The DNA replication timing programme, defined as the temporal order in which genomic loci replicate, is of fundamental importance to maintaining genome stability. In human cells, replication timing is routinely measured using Repli-seq which reveals that the human genome is divided into replication timing domains that replicate in early or late S-phase. These timing domains are associated with gene expression, chromatin compaction, and three-dimensional genome organisation. In particular, late-replicating domains can be hotspots of genomic instability under replication stress. Because of the parasite’s unusual cell cycle, Repli-seq cannot be used in the malaria parasite *Plasmodium falciparum*, leaving the open question as to whether its genome is divided into replication timing domains and how this might impact evolution of the parasite. Here, we leverage the power of Oxford Nanopore Technologies long-read DNA sequencing together with DNAscent to determine, for the first time, the replication timing programme of *P. falciparum*. We show how it relates to elevated rates of DNA breaking and replication fork stalling, and predict that these stalls are predominantly transient.

## Introduction

In eukaryotic cells, the average time in S-phase when each genomic locus replicates is known as DNA replication timing. In human cells, certain regions of the genome replicate reproducibly early or late in S-phase; these are known as replication timing domains. The temporal order in which genomic loci replicate is associated with fundamental features of functional genomics, including gene expression, chromatin compaction, and three-dimensional genome organisation (1). Moreover, DNA replication timing is commonly dysregulated in cancer cells, highlighting its importance to maintaining genome stability (2).

In mammalian cells, the primary assay for DNA replication timing is Repli-seq (3). Briefly, cells are grown in the thymidine analogue bromodeoxyuridine (BrdU) and then sorted into bins using fluorescence-activated cell sorting (FACS). Immunoprecipitation of BrdU-labelled DNA followed by Illumina sequencing allows labelled reads from each bin to be mapped to the genome. At each genomic locus, a weighted average is computed from the number of reads from each bin mapping to that location. This results in a so-called DNA replication timing profile across the genome, taking the form of a curve with chromosomal position along the x-axis and replication timing along the y-axis. Recent variants and expansions upon this technique have included high-resolution Repli-seq (4) and single-cell Repli-seq (5).

The malaria parasite *Plasmodium falciparum* has a highly unusual cell cycle, undergoing a form of asexual multiple fission called schizogony. During the pathogenic blood stage of infection, merozoites infect host erythrocytes and undergo a cell cycle that lasts approximately 48 hours. In stark contrast to human cells, these schizonts perform approximately 4-5 asynchronous rounds of genome replication beginning at approximately 28 hours post-infection (HPI), and thus produce upwards of 16 copies of their genome before cytokinesis takes place (6, 7). These multiple, asynchronous rounds of DNA replication make it impossible to use Repli-seq to determine replication timing. Hence, whether *P. falciparum* has a replication timing programme and, if it does, how it relates to the functional genomics of the parasite are important unanswered questions.

We and others have used Oxford Nanopore Technologies (ONT) long-read DNA sequencing, combined with AI, to measure the replication fork speed across the genomes of *P. falciparum* (8, 9) and *P. knowlesi* (10). Moreover, we have shown that our software, called DNAscent, can measure the frequency of replication fork stalling in human cells (11). Here, we combine DNAscent with mathematical modelling to reveal the replication timing programme of *P. falciparum* and show that both telomeres and key virulence gene families are late-replicating. We then show that replication forks have a very high rate of stalling in *P. falciparum*, but that these stalls are likely to be transient.

## Results

We previously generated high-depth ONT sequencing data from highly synchronised *P. falciparum* parasites at two time-points: 30 HPI (early S-phase) and 36 HPI (mid/late S-phase) (8). To mark the genomic position of moving replication forks, the parasites were sequentially pulsed with EdU (7.5 minutes) and BrdU (7.5 minutes) followed by a thymidine chase prior to parasite harvest. We then used DNAscent v3.0.2 to call the positions of EdU and BrdU on each ONT-sequenced molecule followed by segmentation into replication fork tracks. These data, collected from five ONT R9.4.1 MinION sequencing runs, were then pooled for analysis.

### *Plasmodium falciparum* shows a distinct replication timing programme.

Across the five MinION sequencing runs, this resulted in 1,821 replication fork tracks at the 30 HPI timepoint and 26,006 replication fork tracks at the 36 HPI timepoint. Given the greater amount of replication taking place at 36 HPI, we used this timepoint for all subsequent analysis. We divided the *P. falciparum* genome into 2,337 non-overlapping 10-kilobase (kb) bins. A fork track was assigned to a bin if its 5’ end overlapped that bin, so each track was assigned to exactly one bin. This resulted in 99.7% coverage of the *P. falciparum* genome in fork tracks at an average depth of 11.1 × (Figure 1a). Importantly, depth was fairly uniform across the genome with only 70 bins having a depth lower than 5×.

Following (12), at each 10 kb bin, we computed the replication fork directionality

**Fig. 1.**
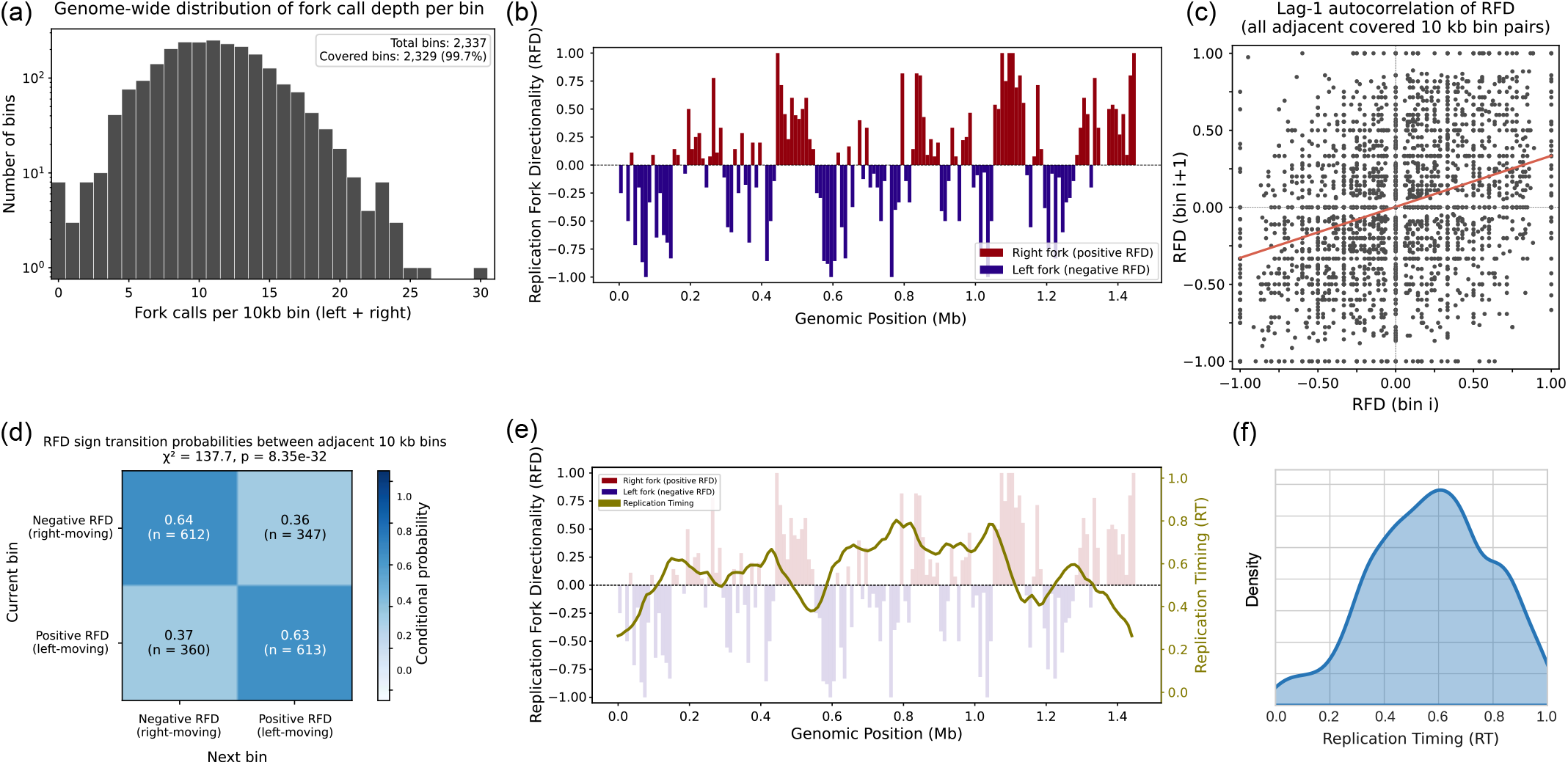
**(a)** Number of DNAscent-called replication fork tracks mapping to each 10-kb bin. Both leftward- and rightward-moving fork calls are included. **(b)** Replication fork directionality across *P. falciparum* chromosome 7, indicating whether each 10-kb bin is predominantly replicated by a rightward- (red) or leftward-moving (blue) fork relative to the 5’ *→* 3’ reference genome assembly. **(c)** Scatter plot of the replication fork directionality of each bin (x-axis) and its immediate neighbour (y-axis) showing a significant positive correlation (Pearson *R* = 0.329; *p <* 0.001; *N* = 2, 315). **(d)** Transition probability matrix indicating the probability of changing RFD while moving across neighbouring bins. These transitions were non-random according to a chi-square goodness-of-fit test (*p* = 8.35 *×* 10^−32^). **(e)** Replication timing curve (gold) overlaid on RFD for *P. falciparum* chromosome 7. Values near 1 indicate regions that replicate early while values near −1 indicate regions that replicate late. **(f)** Density plot of the genome-wide replication timing for *P. falciparum*.

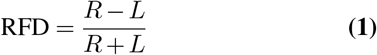

where *R* is the number of rightward-moving forks and *L* is the number of leftward-moving fork tracks mapping to that bin relative to the reference genome written in the 5’ → 3’ direction. It follows that *RFD* is bounded on the closed interval [−1, 1] where *RFD* ≈ −1 indicates that the bin is replicated predominantly by leftward-moving forks and *RFD* ≈ 1 indicates that the bin is replicated predominantly by rightward-moving forks. We found that there were characteristic patterns across the genome where loci were predominantly replicated by forks moving in a particular direction (Figure 1b). Moreover, we showed that these patterns were robust to our choice of a 10 kb bin size (Supplemental Figure S1).

While we achieved high-depth coverage of the *P. falciparum* genome in fork tracks, we sought to confirm that the patterns of RFD that we observed were not the product of random noise. If there was no tendency for a locus to be replicated by a leftward- or rightward-moving fork and patterns of the type shown in Figure 1b were noise, we would expect there to be no correlation between the RFD of a bin and that of its neighbours. We therefore plotted the RFD of adjacent bins and found that they had a significant positive correlation (Figure 1c). Furthermore, we measured the probability that neighbouring bins transition between predominantly left-moving and predominantly right-moving, which was shown to be non-random by a chi-square goodness-of-fit test (Figure 1d). Hence, we conclude that the distinct pattern of RFD observed in Figure 1b, and across the *P. falciparum* genome, is non-random and that the depth shown in Figure 1a is sufficient to reveal this pattern.

As shown in (13), this mean replication time is related to the replication fork directionality by

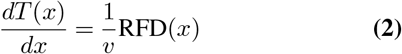

where the functions *T* (*x*) and *RFD*(*x*) are the replication timing and replication fork directionality at bin *x*, respectively, and *v* is the replication fork speed. While we could have measured the replication fork speed in each bin such that *v*(*x*) is a function of position, the depth in fork tracks was still likely to be too small to measure a high-confidence, position-specific fork speed. For robustness, we therefore assumed a constant fork speed of *v* = 0.6 kb/min, noting that replication forks move at a relatively constant speed throughout the genome (8). Solving this differential equation enabled us to convert RFD to a replication timing curve (Figure 1e). Replication timing curves for all *P. falciparum* chromosomes are shown in Supplemental Figure S2.

These curves indicated that *P. falciparum* chromosomes have a non-uniform replication timing, as some regions of the genome replicate early whilst others replicate late. A replication timing programme has never previously been demonstrated outside model eukaryotes with binary-fission cell cycles, such as human cells or *S. cerevisiae* (1); nor has replication timing ever before been derived from DNAscent data alone. Doing so has allowed us to identify clear differences between *P. falciparum* and human cells. The replication timing of human cells tends to be bimodal with peaks at early and late S-phase (14). In *P. falciparum*, we instead observed a unimodal rather than a bimodal pattern (Figure 1f).

### Key chromosomal features and gene families replicate characteristically early or late

The replication timing curves suggested that telomeres may replicate late whilst centromeres may replicate early (Figure 2a). Hence, we sought to determine whether this was the case across the *P. falciparum* genome. We annotated each chromosome with the location of its centromere, as defined by CENH3-covered regions of the chromosome (15). Doing so revealed that centromeres replicate early across all 14 *P. falciparum* chromosomes, bringing *P. falciparum* in line with other eukaryotes including budding yeast (16), fruit flies (17), and mice (18) (Figure 2b). However, this contrasts with the human genome where centromeres tend to replicate in mid-to-late S-phase (19). We used long read assemblies of the *P. falciparum* lab-adapted clone 3D7 to delineate the boundaries of subtelomeric regions for each chromosome (20). Doing so indicated that subtelomeres replicate late across all 14 chromosomes (Figure 2b). This aligns with budding yeast (16), but again puts *P. falciparum* at odds with human cells where telomeres replicate throughout S-phase (21, 22).

**Fig. 2.**
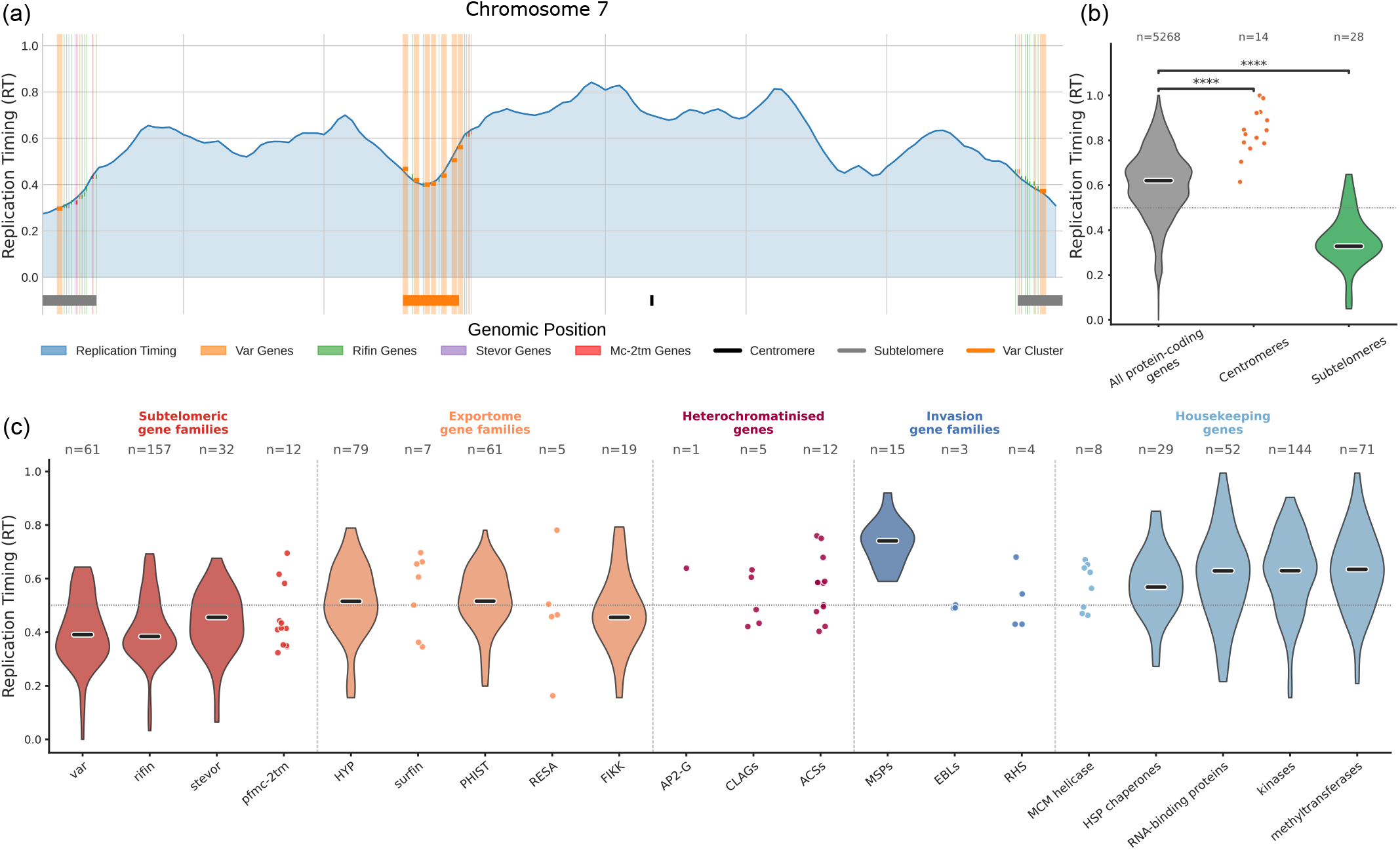
**(a)** The replication timing curve from Figure 1e, annotated with *var, stevor, rifin*, and *pfmc-2tm* genes as well as the centromere and subtelomeres. **(b)** The replication timing of all protein-coding genes compared to the replication timing of the centromere on each chromosome (orange) and subtelomeres (green). Due to a small number of points, centromeres are shown as individual points rather than a violin plot. Results are from a two-sided Mann-Whitney U test (****: *p <* 0.0001). **(c)** Replication timing of key gene families, excluding pseudogenes. Where the number of genes is low, distributions are shown as individual points rather than violin plots.

Again prompted by the annotated replication timing curve in Figure 2a, we next investigated the replication timing of key gene families. *P. falciparum* has several families of variantly expressed virulence genes, many of which encode variant surface antigens. These genes play important roles in pathogenesis and immune evasion (23). They are often, but not always, encoded at sub-telomeric locations and tend to be physically clustered and heterochromatinised within the nucleus (24, 25). Hence, we hypothesised that they would replicate late. In particular, we investigated the family of approximately 60 *var* genes that encode the virulence factor PfEMP1, which is key for mediating cytoadherence of infected erythrocytes to endothelial receptors (26–28); *rifin* (repetitive interspersed family) genes, which encode proteins with emerging roles in immune evasion via mimicry-based engagement of leukocyte receptors (29–33); *stevor* (subtelomeric variable open reading frame) genes, which may have a role in rosetting (34, 35); and the little-characterised *pfmc-2tm* (*P. falciparum* Maurer’s clefts 2 transmembrane) family of genes (36, 37). Each of these gene families undergoes some level of expression switching by continually changing which gene or genes are transcriptionally active, thereby maintaining antigenic variation and immune evasion. In the case of *var* genes, expression switching is controlled epigenetically by histone methylation, acetylation and heterochromatin protein 1, with the active gene in a euchromatic state and silent genes in condensed heterochromatin (38, 39). Indeed, we determined that all four gene families replicate late (Figure 2c). This timing appeared to be determined primarily by chromosomal location. Other exportome gene families replicated only slightly late, consistent with their being less proximal to telomeres, and heterochromatinisation per se was not sufficient to impose late replication because genes such as *ap2-g* and *clag*, which are silenced by non-telomeric islands of heterochromatin (40), did not necessarily replicate late (Figure 2c).

### Replication timing is associated with gene expression and chromatin state

In human cells, DNA replication timing is closely associated with both chromatin compaction and gene expression, with early-replicating regions of the genome associated with open chromatin and highly transcribed genes (41). Moreover, replication timing is associated with nuclear organisation of the genome which can impact genomic rearrangements. However, genome organisation in *P. falciparum* is unusual compared to human cells: the majority of chromatin is open and loosely transcribed, while islands of heterochromatin are limited to variantly expressed genes such as *var* genes and *ap2-g* (40). Hence, we sought to determine whether the relationship between chromatin state, gene expression, and replication timing holds in *P. falciparum*.

To determine whether replication timing was associated with chromatin state, we compared replication timing to ATAC-seq from (42). There was a general association between early-replicating regions and open chromatin visible on single chromosomes (Figure 3a). Genome-wide, there was a slight but significant correlation (Spearman *ρ* = 0.126; *p <* 0.001; *N* = 2, 337) between replication timing and chromatin accessibility as measured by ATAC-seq (Figure 3b). Hence, in *P. falciparum*, replication timing is associated with chromatin accessibility, albeit with a lower correlation than is observed in human cells (3).

**Fig. 3.**
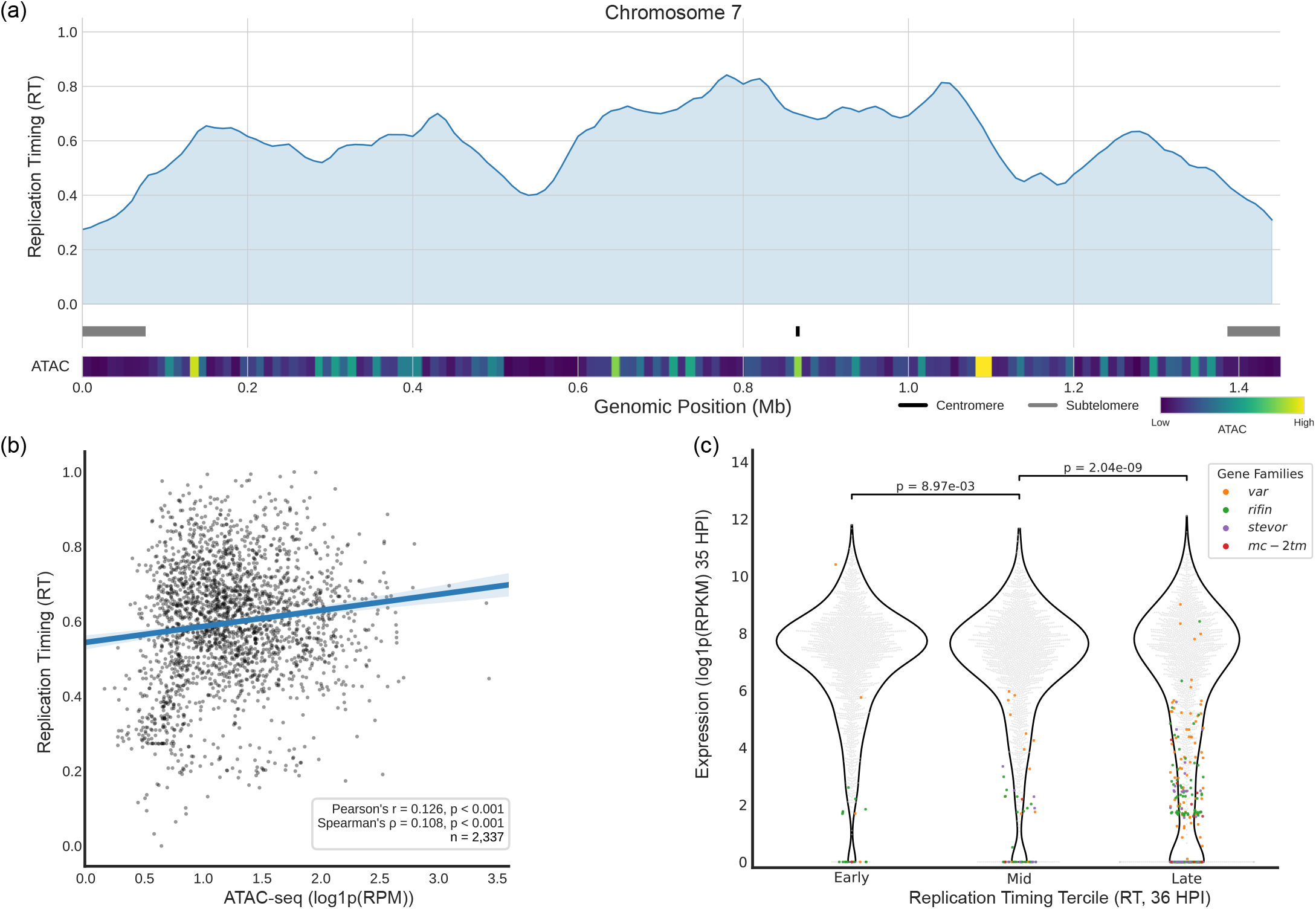
**(a)** Replication timing of chromosome 7 with ATAC-seq data. Lighter values indicate loci of more accessible chromatin. **(b)** Correlation of replication timing with ATAC-seq, represented as log1p(RPM). **(c)** The genome was divided into terciles by replication timing, and the expression of genes in each tercile is plotted as log1p(RPKM). Points corresponding to members of four antigenic variation gene families are shaded. P-values represent the results of a one-sided Mann-Whitney U test against the alternative hypothesis that gene expression decreases from early to late replication timing.

To determine whether gene expression followed the same trend, we partitioned the genome into early-, mid-, and late-replicating terciles based on replication timing and examined the expression of genes in each tercile. There was a clear and statistically significant decrease in gene expression moving from early-replicating to late-replicating terciles (Figure 3c). However, as with chromatin accessibility, the association with replication timing was relatively modest, and may be substantially due to the late-replicating tercile encompassing subtelomeric regions, which include gene families like *var* where the majority of the family is silenced.

Together, these data indicate that there is an association between chromatin openness, actively transcribed genes, and early replication timing. Although these trends are not as strong as those observed in human cells, the weaker association is consistent with the predominantly euchromatic *P. falciparum* genome, which is only punctuated by relatively small regions of heterochromatin (40). This also serves as a potential explanation for the lack of bimodality in replication timing observed in Figure 1f. Moreover, as exemplified by *ap2-g* and *clag* genes in Figure 2c, replication timing is associated with, but not imposed by, whether a locus is euchromatic or heterochromatic.

### Replication fork speed is consistent across S-phase but changes at key gene families and genomic features

We and others have shown that replication fork speed is slower in early-replicating regions of the human genome (11, 43), while the replication fork speed in budding yeast has been shown to be remarkably consistent across S-phase (44). Therefore, we sought to determine whether replication fork speed in *P. falciparum* was variable or constant with respect to replication timing. We were particularly interested in whether the *P. falciparum* genome contained loci that were inherently difficult to replicate, mirroring common fragile sites in human cells.

When considering replication fork speed across replication timing terciles, we did not observe an increase in fork speed as timing terciles moved from early to late (Figure 4a). However, we observed a statistically significant decrease in replication fork speed as forks moved through centromeres and telomeres (Figure 4b) which is consistent with the findings in (9). This contrasted with a significant increase in fork speed as forks moved through the *stevor* and *pfmc-2tm* families. There was also evidence to suggest that fork speed decreased at *var* genes and increased at *rifin* genes, though these did not reach statistical significance (*p* = 0.055 and *p* = 0.077, respectively).

**Fig. 4.**
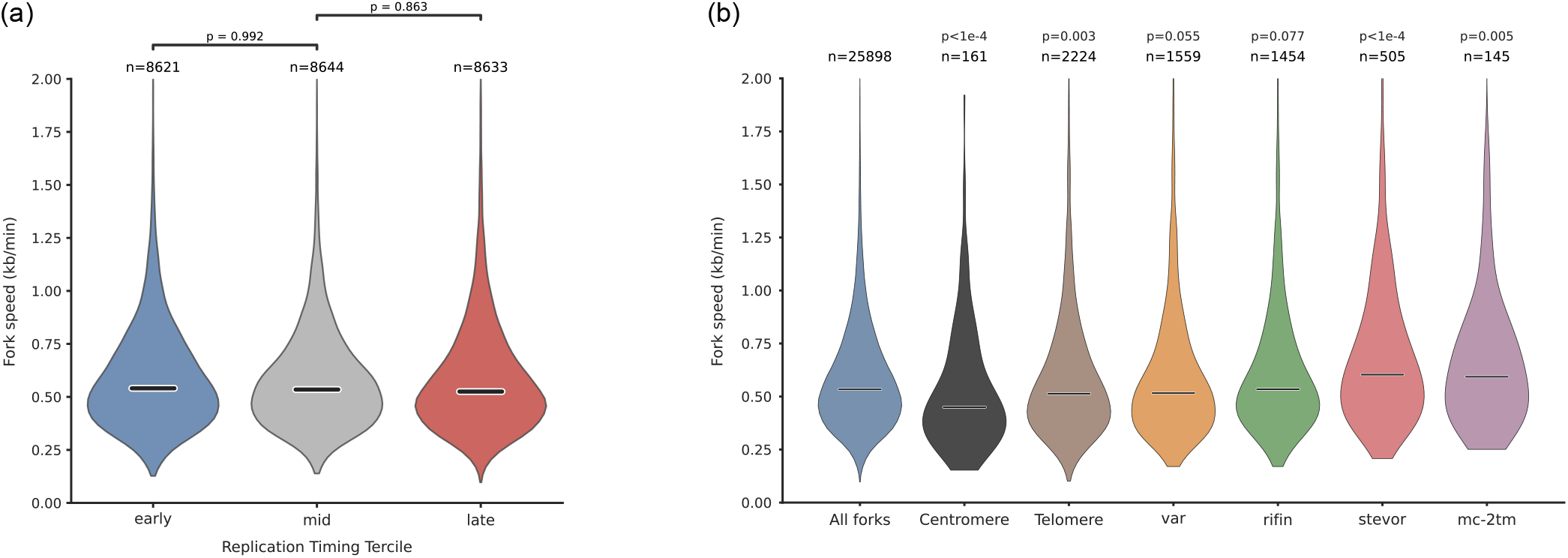
**(a)** Replication fork speed, as measured by DNAscent, in regions of the genome corresponding to each replication timing tercile. P-values represent the results of a one-sided Mann-Whitney U test against the alternative hypothesis that replication fork speed increases from early to late replication timing. **(b)** The DNAscent-measured replication fork speed using tracks that mapped to centromeres, telomeres, or the *var, rifin, stevor*, or *pfmc-2tm* gene families. P-values show the results of a two-sided Mann-Whitney U test comparing fork tracks that mapped to each genomic feature to all fork calls across the genome.

While we observed distinct patterns of replication timing for centromeres, telomeres, and the four antigenic variation gene families, we found that replication fork speed does not vary according to replication timing alone. This suggests that attributes of these specific genomic features, rather than the time at which they replicate, are responsible for the changes observed in replication fork speed.

### *Plasmodium falciparum* shows elevated levels of fork stalling compared to human cells

We previously used DNAscent to identify replication fork stalling, which manifests as a sharp drop-off in BrdU incorporation created when a fork stops moving during the BrdU pulse. DNAscent measures the steepness of this drop-off to create a stall score, where fork tracks with a stall score near 0 indicate a fork that is unlikely to have stalled, while a stall score near 1 indicates a fork that is very likely to have stalled. In budding yeast, DNAscent was able to identify fork stalls at replication fork barriers in rDNA (45, 46). We observed the same effect in human cancer cells when treated with inhibitors that target the DNA replication stress response pathway. However, in untreated, non-cancerous human RPE1 cells, we observed negligible levels of stalling in agreement with expectation (11). We applied this method to *P. falciparum* and observed remarkably high stall scores at both 30 and 36 HPI (Figure 5a). It is noteworthy that these high stall scores were observed in the absence of treatment with any anti-malarial agents or inhibitors. Hypothesising that the high level of AT-richness in *P. falciparum* was contributing to these high rates of stalling, we measured the stall scores in *P. knowlesi*, which has a more moderate 61% AT-richness. We found that the rate of fork stalling in *P. knowlesi* was elevated and significantly higher than that of human RPE1 cells, and that *P. falciparum*, in turn, had a higher rate of stalling than *P. knowlesi* (Figure 5a).

**Fig. 5.**
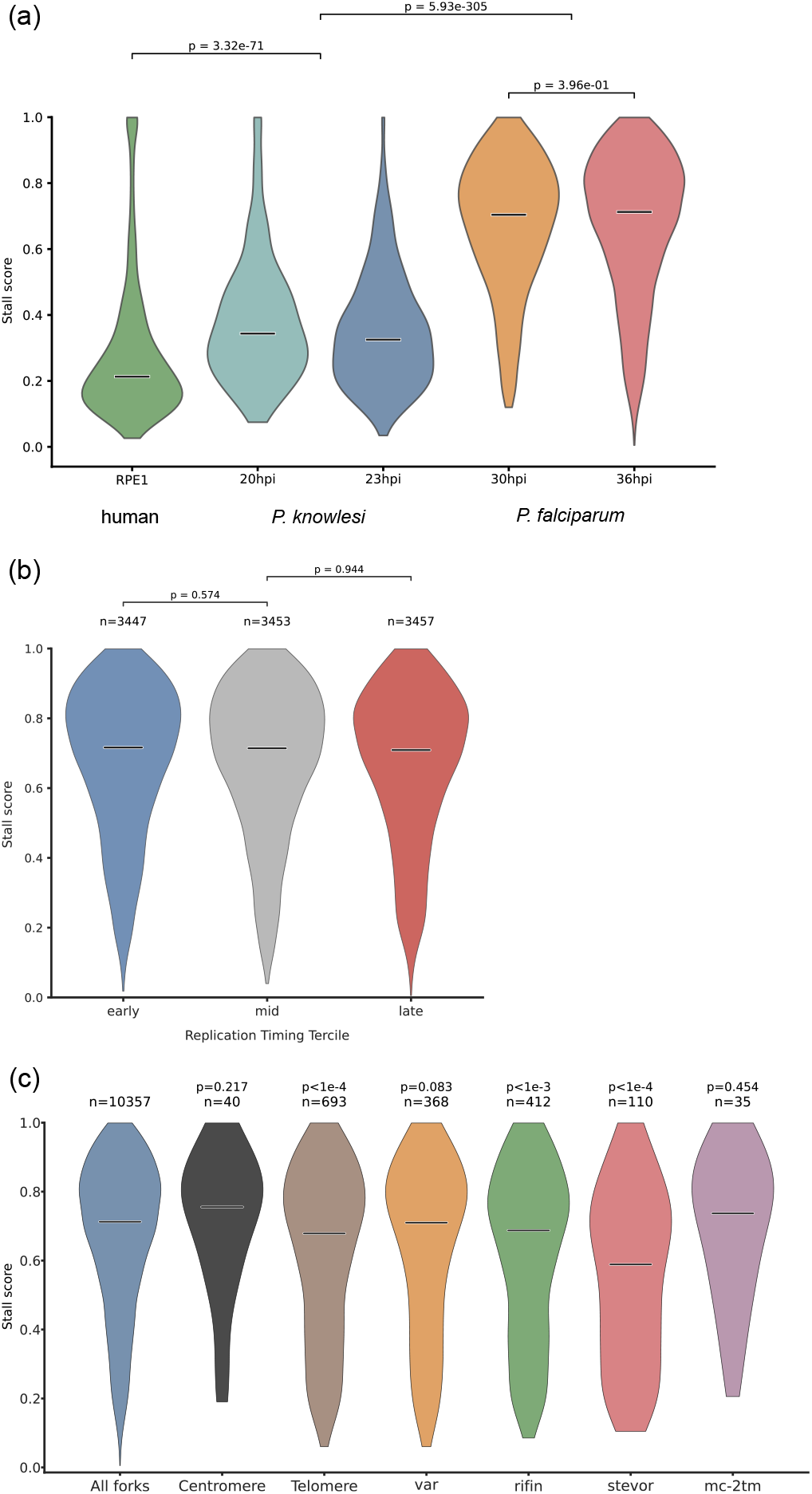
**(a)** Distribution of stall scores from all replication fork calls across the genome of human RPE1 cells, *P. knowlesi*, and *P. falciparum*. Horizontal bars indicate the median. P-values indicate the results of a two-sided Mann-Whitney U test. When comparing between two different species where two timepoints post-infection were available, stall scores from both timepoints were pooled. **(b)** Stall scores separated into replication timing terciles, as in Figures 3c and 4a. P-values represent the results of a one-sided Mann-Whitney U test against the alternative hypothesis that replication fork stalling decreases from early to late replication timing. **(c)** Stall scores by genomic feature, mirroring the fork speed analysis in Figure 4b. P-values show the results of a two-sided Mann-Whitney U test comparing fork tracks that mapped to each genomic feature to all fork calls across the genome.

Previously, we observed a decrease in replication fork stalling over S-phase in HCT116 cells (11). However, as with replication fork speed, we did not observe this trend in *P. falciparum* (Figure 5b), whereas there was a significantly lower rate of stalling at telomeres and at *rifin* and *stevor* genes. Interestingly, there was some evidence for elevated rates of stalling at centromeres, but this did not reach statistical significance (Figure 5c).

Taken together, these observations indicate that the elevated fork stalling in *P. falciparum* is a genome-wide phenomenon rather than one confined to particular loci or replication timing domains. The absence of a clear trend across replication timing terciles, together with the reduced stalling at telomeres and at the *rifin* and *stevor* gene families, suggests that fork stalling in *P. falciparum* is driven principally by local sequence composition and specialised chromatin rather than by bulk chromatin accessibility or the timing of replication in S-phase.

### DNA breaks at a high rate during *P. falciparum* replication

DNAscent, as of v4.1.1, can measure the levels of DNA breaking during S-phase from DNA sequenced on both R9.4.1 and R10.4.1 flow cells. If DNA breaks at a replication fork during the analogue pulse, this will result in the analogue track running to the read end. However, an analogue track running to the read end can occur due to a DNA break or due to random fragmentation of the sequencing library, and the frequency of the latter depends on the replication fork speed, duration of the analogue pulse, and quality of the library preparation. The subprogram DNAscent seeBreaks normalises for these three factors, allowing us to measure whether more analogue tracks run to the read end than would be expected by chance.

In *P. falciparum*, DNAscent seeBreaks called elevated breaking in four out of the five 36 HPI replicates (Figure 6a). As before, we partitioned the *P. falciparum* genome into replication timing terciles and examined the level of breaking over replication. At the level of individual replicates, levels of breaking were elevated but did not show a statistically significant change across replication timing terciles, nor did we observe a change across timing when replicates were pooled at each tercile, although there was a near-significant trend towards lower breakage at later timepoints (Figure 6b).

**Fig. 6.**
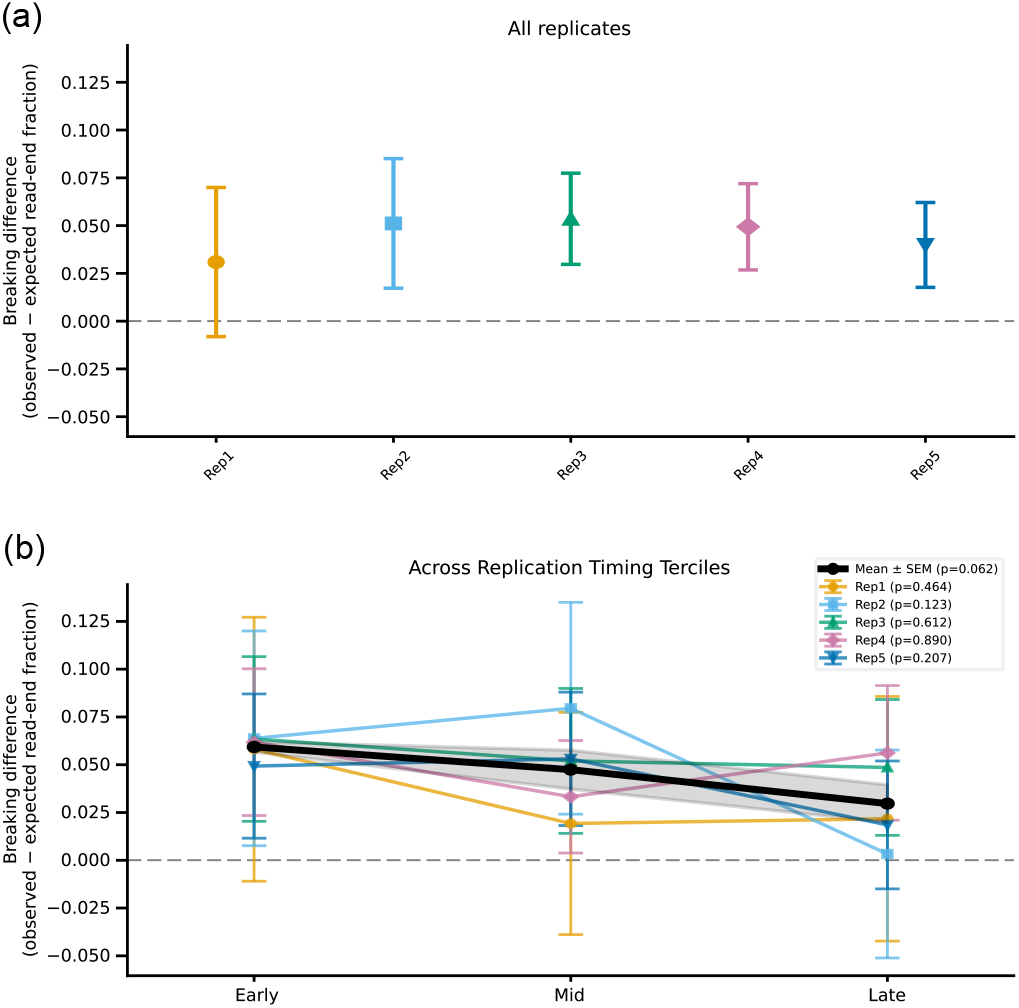
**(a)** The results of DNAscent seeBreaks at 36 HPI across five biological replicates. Markers indicate the mean difference between the observed and expected number of fork tracks at read-ends such that values above 0 are indicative of more observed tracks than would be expected by chance. Whiskers indicate the 95% confidence interval for the mean. **(b)** The results of each replicate partitioned into replication timing terciles. Markers indicate the mean difference between the observed and expected number of fork tracks at read-ends at each tercile. Whiskers indicate the 95% confidence interval for the mean. P-values are the result of a two-sided weighted linear trend test with weights *w*_*i*_ = 1*/SE*^2^ for each tercile *i*. Black points indicate the results of pooling replicate means at each timing tercile, and the shaded area indicates the standard error of the mean. Per-replicate trend z-scores were combined across the five replicates using Stouffer’s method 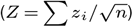 giving the pooled two-sided p-value shown.

### Mathematical modelling predicts that replication fork stalling is transient

Given the high rates of replication fork stalling we observed in *P. falciparum*, we asked what the implications might be for the timely completion of genome replication. *P. falciparum* genome replication has been estimated to complete in approximately 60 minutes (6, 7), which is not particularly long compared to other organisms with a similar genome size. To achieve this timely completion despite very frequent fork stalling, we would expect frequent replication origin firing and therefore a small inter-origin distance. Estimates of *P. falciparum* inter-origin distance (IOD) from DNA fibre labelling are approximately 60-70 kb (48) which, while shorter than in human cells (49), did not appear low enough to reflect such elevated levels of fork stalling. Furthermore, DNA breaking, while elevated, was also not elevated to the extent that we would have expected for the elevated stalling that we observed.

We therefore created a mathematical model, fitted to the *P. falciparum* 36 HPI replication timing programme, using the procedure from (47), which produced a close fit to the data (Figure 7a). We then created two variants of this model: one where fork stalls were transient and resumed quickly relative to the duration of the pulse, resulting in a slow effective fork speed, and another where fork stalling was permanent (Figure 7b). Both models were able to fit the replication timing data (Figure 7c), although the model with permanent stalling was slower to converge. Moreover, the curve indicating the rate of origin firing also took the same shape across the genome in both models although, as expected, the origin firing rate was fit to a higher value in the model with permanent fork stalling (Figure 7d).

**Fig. 7.**
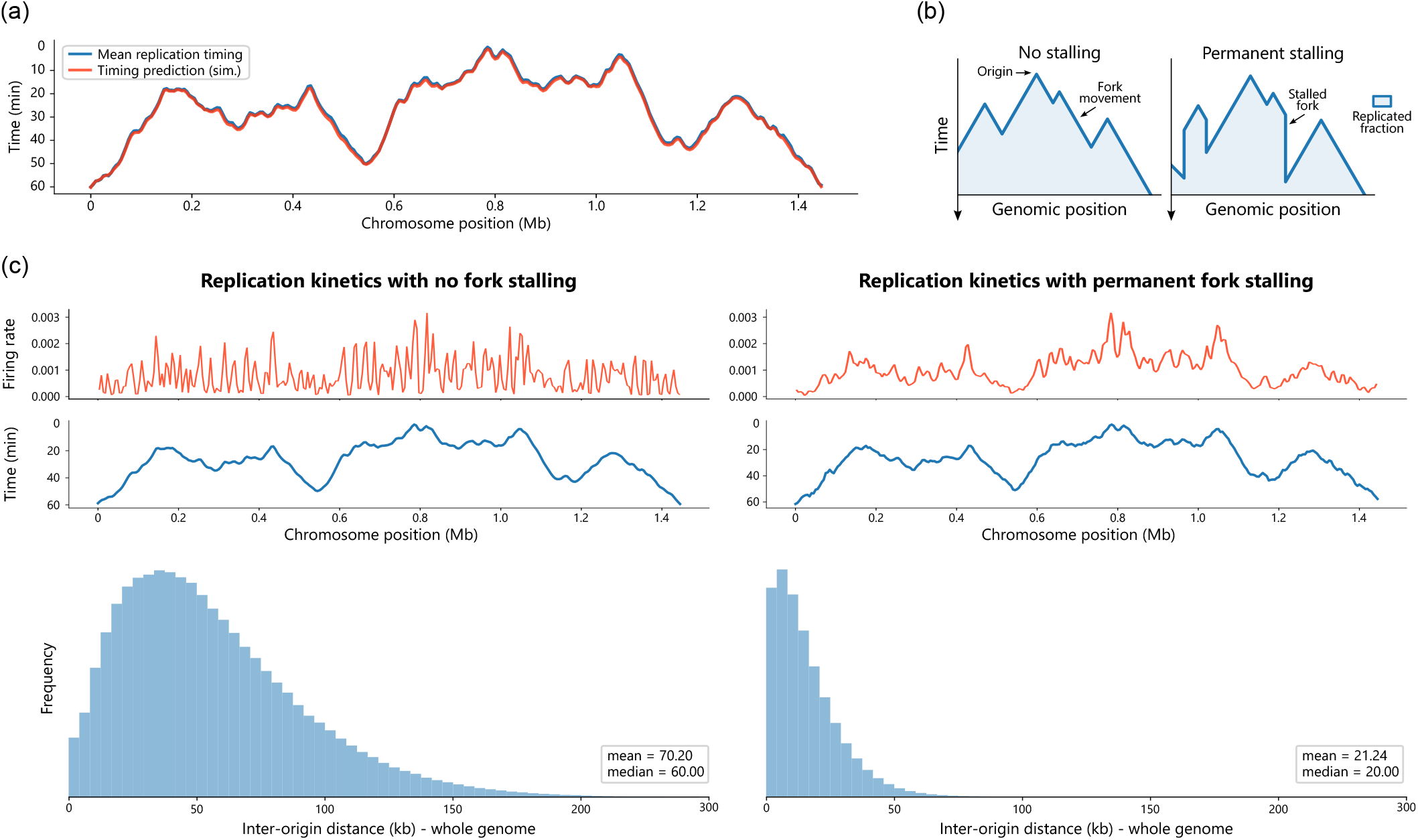
**(a)** The simulated replication timing curve (red) from a mathematical model fit to replication timing data (blue) using the methodology from (47), shown here for *P. falciparum* chromosome 7. Replication fork stalling is not included in the model and the simulated curve is averaged over 10,000 simulations. **(b)** A variant of the model with permanent replication fork stalling included. The impact of replication fork stalling is shown on the replication timing curve of a single simulation. **(c)** The origin firing rate (upper), replication timing curve (middle), and distribution of inter-origin distances (lower) for a mathematical model fit to replication timing data when the model does (right) and does not (left) include replication fork stalling. Model results are averaged over 10,000 simulations. Firing rate curves and replication timing curves are shown for *P. falciparum* chromosome 7 while the distribution of inter-origin distances is taken over the whole genome.

Measuring the inter-origin distance from the model with transient stalling revealed a mean IOD of 70 kb that is consistent with actual data from DNA fibre labelling (48). However, the model with permanent stalling had a mean IOD of only 21 kb, which is in sharp contrast to estimates from DNA fibre. Moreover, if the inter-origin distance was only 21 kb, we would expect to see multiple origins frequently on our ONT reads given that the mean length of reads that included a fork call was 37.4 kb (8). However, of the 3,018 reads that included an origin call, only 167 of them included more than one origin call. Hence, we conclude that fork stalls, while frequent in the *P. falciparum* genome, are usually transient and recoverable pauses in replisome progression, presumably as replication forks navigate the highly AT-rich *P. falciparum* genome.

## Discussion

Here, we have shown that replication fork directionality, as determined by single-molecule analysis with DNAscent, can be mapped to a replication timing curve in the malaria parasite *P. falciparum*. Erythrocytic schizogony means that more traditional assays for measuring DNA replication timing, such as Repli-seq, are intractable, and hence our method has revealed the replication timing programme of *P. falciparum* for the first time. Using this replication timing information, we have shown that genomic features such as telomeres, centromeres, and antigenic variation genes replicate at characteristic times, and that replication timing is associated with gene expression and chromatin openness. We further showed that *P. falciparum* has highly elevated levels of fork stalling compared to both *P. knowlesi* and human RPE1 cells, but that most of these stalling events are likely to be transient pauses rather than permanent stops, enabling the timely completion of genome replication.

While we did not observe significant trends between replication fork speed, replication fork stalling, DNA breaking, and replication timing, we did observe markedly different behaviour at specific genomic features that replicated characteristically early or late. There was some evidence for replication stress at centromeres: forks markedly slowed as they moved through them, and stalling appeared elevated, although this did not reach statistical significance. Telomeres also showed replication fork slowing, albeit not to the same extent as centromeres, but conversely to centromeres, they showed less fork stalling than the rest of the genome. In *P. falciparum*, centromeres are low-complexity higher-order repeats packaged in CENH3-containing kinetochore chromatin (15, 50), while telomeres comprise G-quadruplex-forming GGGTT(T/C)A repeats bound by dedicated telomeric proteins (51, 52). Notably, telomeres are not AT-rich, yet they still slowed the replisome, and we have previously shown that bulk fork speed is unchanged even in regions exceeding 90% AT content (10). Together, this suggests that the *P. falciparum* replisome is able to robustly navigate extreme AT-content without slowing, as we demonstrated by comparing replication fork speeds in *P. falciparum, P. knowlesi*, and human cells (10). Instead, fork slowing at centromeres and telomeres is more likely driven by their repetitive structure and specialised chromatin. The AT-richness of the *P. falciparum* genome is instead associated with genome-wide fork pausing (Figure 5) which, because it is largely recoverable, is part of the slow effective fork speed that we measured.

Importantly, it appears that the speed and stalling of replication forks are interrelated but distinct phenomena, which can have distinct biological consequences. We observed a significant increase in replication fork speed in *stevor* and *pfmc-2tm* genes, with no significant change at genes in the *var* and *rifin* families. We also observed a significant decrease in stalling at *rifin* and *stevor* genes, with no significant change in *var* or *pfmc-2tm* genes. It is noteworthy that the primary sequences of *rifin, stevor*, and *pfmc-2tm* genes are less repetitive than the genome-wide average (Supplemental Figure S3) (53). The higher complexity sequence, together with these genes being largely transcriptionally silent, is consistent with a general decrease in replication stress. It is also noteworthy that *pfmc-2tm* genes are distinguished from *rifin* and *stevor* by very high, rather than low, AT-richness. This may explain why rates of stalling in *pfmc-2tm* were higher compared to these other two families. The complexity of *var* genes is much closer to the genome-wide average than *stevor, rifin*, or *pfmc-2tm*, and correspondingly, we did not observe a significant difference in either speed or stalling as replication forks moved across *var* genes.

While DNAscent has a high spatial resolution, it measures an effective fork speed: how far a replication fork moved during the analogue pulse, with relatively little information given about the dynamics of the replication fork during the pulse. Those dynamics can only be revealed if there are high rates of pausing at the end of the pulse, which manifest as high stall scores. In fact, transient stalling has probably occurred throughout the pulse, resulting in a low effective fork speed. The speed and stalling of replication forks measured on single molecules using analogue pulse-chase must therefore be interpreted with caution, as fork behaviour can be masked in a way that depends on the fork speed and the duration of the analogue pulses.

The uniquely high rates of replication fork stalling that we detected in *P. falciparum* seemed to result from mainly transient pauses, but this nonetheless underscores that *P. falciparum* is under high replication stress during erythrocytic schizogony, especially when taken together with the slow replication fork speed. This raises the question of what the purpose of this replication stress might be, if indeed there is one. It may be a detrimental, but ultimately worthwhile, byproduct of a highly AT-rich genome that drives adaptive genome evolution via microstructural plasticity (54). Alternatively, it may be synergistic to driving adaptive genome evolution by facilitating frequent DNA breaks and copy number changes (55, 56). Prime candidates for fast-evolving regions are the antigenic variation gene families, although we have demonstrated that *rifin* and *stevor* genes appear under less replication stress than the genome-wide average. However, it may be the case that all gene families are not created equally in this regard. Protecting *rifin* genes from replication stress may be evolutionarily advantageous to preserve their LILRB binding properties (57). On the other hand, *var* genes exhibited levels of replication stress near the genome-wide average and mutation may be more advantageous in this gene family. Indeed, *var* genes were shown to be particularly recombinogenic when replication stress was induced via knockout of RecQ helicase (58). Ultimately, it remains an open question whether the pervasive replication stress we observed in *P. falciparum* is a functional feature of parasite biology or simply an unavoidable cost of maintaining an extremely AT-rich genome. Disentangling these possibilities will require future work that directly links replication stress to downstream consequences for genome stability, antigenic variation, and parasite fitness. Determining the role, if any, that this replication stress plays in the evolution and pathogenesis of *P. falciparum* is therefore a promising avenue for future investigation.

## Methods

### Genome-wide map of replication fork directionality

DNAscent-called replication fork tracks at 36 HPI were taken from (8) which are available at NCBI GEO under accession number GSE210010. These fork tracks were called using DNAscent v3.0.2 (https://github.com/MBoemo/DNAscent/releases#release-3.0.2). Coordinates were relative to the *P. falciparum* 3D7 ASM276v2 reassembly (https://www.ncbi.nlm.nih.gov/assembly/GCF_000002765.4/). Left- and right-moving fork calls were pooled across five biological replicates for the 36 HPI timepoint. For each of the 14 nuclear chromosomes, the *P. falciparum* genome was divided into 10 kb bins, and a fork track was assigned to the bin overlapping the 5’ end of the fork call such that each fork call was assigned to a unique bin. RFD was then calculated per bin using Equation 1. Bins containing no fork calls were assigned an RFD of 0 for plotting but excluded from further downstream analyses.

### Replication timing curve

A replication timing curve was calculated for each chromosome via the numerical integration of Equation 2 along each chromosome using the cumulative trapezoidal rule, with the bin midpoint taken as the integration variable and the integral initialised to zero at the first bin. Integrals were computed independently per chromosome, and a genome-wide min-max normalisation was applied across all pooled chromosomes to rescale the replication timing curve to the interval [0, 1]. Under this scheme, the earliest replicating position in the genome was given a value of 0 and the latest position was given a value of 1. This final replication timing value was taken to be

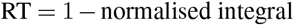

so that values near 1 indicated positions that replicated early and values near 0 indicated positions that replicated late.

### Processing of RNA-seq and ATAC-seq

Strand-specific normalised RNA-seq in bedgraph format was downloaded from NCBI GEO accession GSM2789037 and ATAC-seq in bedgraph format was downloaded from NCBI GEO accession GSM2789027 (42). For the RNA-seq data, length-normalised average signals were calculated while maintaining strand specificity to minimise potential interference from antisense transcription, and the resulting values were scaled to generate the equivalent of RPKM. The resulting values in RPKM (for RNA-seq) and RPM (for ATAC-seq) were subsequently log1p transformed.

### Replication fork stalling

As of DNAscent v3.1.2 (https://github.com/MBoemo/DNAscent/releases#release-3.1.2), DNAscent detects replication fork stalls in data sequenced on R9.4.1 Oxford Nanopore Technologies flow cells (11). The bam files from (8) that were used as inputs to DNAscent v3.0.2 were used as inputs to DNAscent v3.1.2 so that stall scores could be generated for each fork track by DNAscent forkSense. As shown in (11), DNAscent forkSense produces a stall score in the interval [0, 1] for each valid fork track, where scores near 1 indicate that the fork is likely to have stalled during the BrdU pulse. DNAscent forkSense assigns negative integer flags to any invalid fork track to indicate tracks that end in a termination site, run off the read, or are proximal to an indel greater than 100 bp in length. Fork tracks with these negative integer flags were excluded from the analysis.

### Breaking during replication

DNA breaking during replication was measured on all five *P. falciparum* 36 HPI biological replicates using DNAscent seeBreaks v4.1.1 (https://github.com/MBoemo/DNAscent/releases#release-4.1.1) using the DNAscent detect output and fork calls from DNAscent v3.1.2. Fork calls and DNAscent detect outputs were then split into replication timing terciles for each biological replicate. DNAscent seeBreaks was then run separately on each timing tercile for each biological replicate, or on each timing tercile with biological replicates pooled across that tercile.

### Mathematical modelling (no fork stalling)

The prediction of inter-origin distances uses the methodology introduced in (47). Each chromosome was discretised to one-kilobase resolution and every site *j* was treated as a potential origin of replication with an intrinsic firing rate *f*_*j*_. The firing time of the origin at site *j* was modelled as an exponential random variable with rate *f*_*j*_ and fork speed was constant at *v* = 0.6 kb/min. Under these assumptions, the replication time *T*_*j*_ of genomic position *j* was defined as the minimum, over all potential origins *i*, of the firing time at *i* plus the deterministic travel time |*i* −*j*| */v* required for a fork to reach *j*. A closed-form expression for the expected replication time is given by

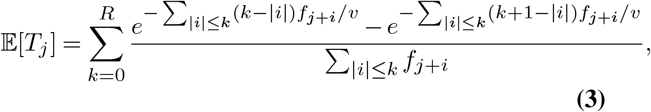

where *R* = 2000 kb is a finite radius of influence determining the distance within which neighbouring origins are allowed to affect the timing of a focal site (47). Given a genome-wide timing profile *T*_*j*_, an initial estimate of the firing-rate landscape was constructed using an analytical approximation valid under uniform firing, after which each *f*_*j*_ was iteratively updated to reduce the discrepancy between the predicted timing E[*T*_*j*_] and the measured profile. After fitting the firing-rate landscape in this way, we simulated the stochastic replication process using the inferred *f*_*j*_ values to simulate a population of 10,000 S-phase cells. For each simulation, we recorded the positions of fired origins and calculated the spacing between neighbouring fired origins.

In practice, directly estimating the derivative of experimental RFD profiles is unstable because pointwise slopes are strongly affected by experimental and computational noise (59). To obtain a stable timing estimate, we integrated Equation 2 over sliding windows of length *l* = 10 kb, which yielded

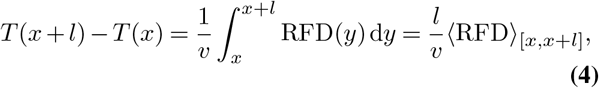

Where ⟨·⟩ _[*x,x*+*l*]_ denotes the average RFD in the interval [*x, x* + *l*]. This smoothing reduced resolution but substantially improved the numerical stability of the inferred timing profile.

The resulting timing curve was then rescaled to match the one-hour duration of S phase in *P. falciparum*. Both models considered origin firing times as a shifted exponential distribution

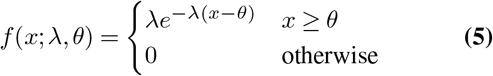

where *θ* = 70 avoided the artificial clustering of early-firing events that would otherwise arise when timing is scaled from zero.

Given the monotonic dependence of the expected replication time E[*T*_*j*_] on the firing rates {*f*_*j*_}, we estimated {*f*_*j*_} by iteratively adjusting each value to reduce the discrepancy between the predicted timing 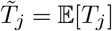 and the inferred timing *T*_*j*_. Following (47), we initialised

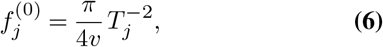

and refined by

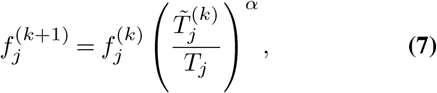

where *α* controlled the update strength. At each iteration, 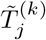 was obtained from Equation 3 with the current firing rates. This fixed-point procedure exploits the monotonic structure of Equation 3, and converged rapidly without requiring large-scale simulations (for *α* = 0.5).

## Mathematical modelling (permanent fork stalling)

To estimate the rate of stalling, we considered the fraction of fork calls with a high stall score (>0.8) and excluded forks with negative integer flags assigned by DNAscent forkSense. This led to 3,398 of 10,357 (32.8%) forks being classified as stalled. The EdU/BrdU labelling period had a total duration of 15 minutes (8) and, using our previously calculated replication fork speed of *v* = 0.6 kb/min, a continuously progressing fork would traverse approximately *L* = 0.6 × 15 = 9 kb. The probability that a fork stalls is therefore

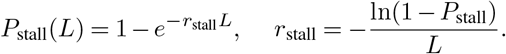

Using *P*_stall_ = 0.328 gives

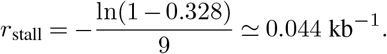

When permanent stalling was included in the model, replication timing depended not only on firing rates but also on the spatial stall rate *r*_stall_. This coupling broke the analytic structure used to obtain the initial guess Equation 6, since the relation between *T*_*j*_ and *f*_*j*_ was no longer directly characterised by Equation 3. As a result, we did not apply the analytic initialisation and instead started from a uniform firing-rate profile. Despite the lack of a closed-form first guess, the multiplicative update Equation 7 remained applicable. In practice, using a smaller update exponent (*α* = 0.2) yielded stable convergence of the fixed-point iterations even in the presence of stalling.

## ACKNOWLEDGEMENTS

This work was supported by the Leverhulme Trust (RPG-2022-028), the Medical Research Council (MR/W031442/1), and the Biotechnology and Biological Sciences Research Council (UKRI1912). KS was funded by a summer research studentship by the British Division of the International Academy of Pathology, St John’s College Cambridge, and the University of Cambridge Department of Pathology. RBL was funded by a PhD studentship provided by the University of Cambridge Department of Pathology (John Lucas Walker Fund) and Trinity Hall Cambridge. SEC was funded by a PhD studentship provided by the University of Cambridge Department of Pathology (Centenary Fund). FB was funded by the Biological Sciences Research Council (UKRI1912) and further supported by a College Research Associateship to Emmanuel College Cambridge. CJM was funded by the European Research Council (ERC) under the European Union’s Horizon 2020 research and innovation programme (ERC-2016-COG 725126). This work was performed using resources provided by the Cambridge Service for Data Driven Discovery (CSD3), operated by the University of Cambridge Research Computing Service (https://www.csd3.cam.ac.uk), and supported by Dell EMC and Intel through Tier-2 funding from the Engineering and Physical Sciences Research Council (capital grant EP/T022159/1) and DiRAC funding from the Science and Technology Facilities Research Council (https://dirac.ac.uk).

## Supplementary Material

**Fig. S1.**
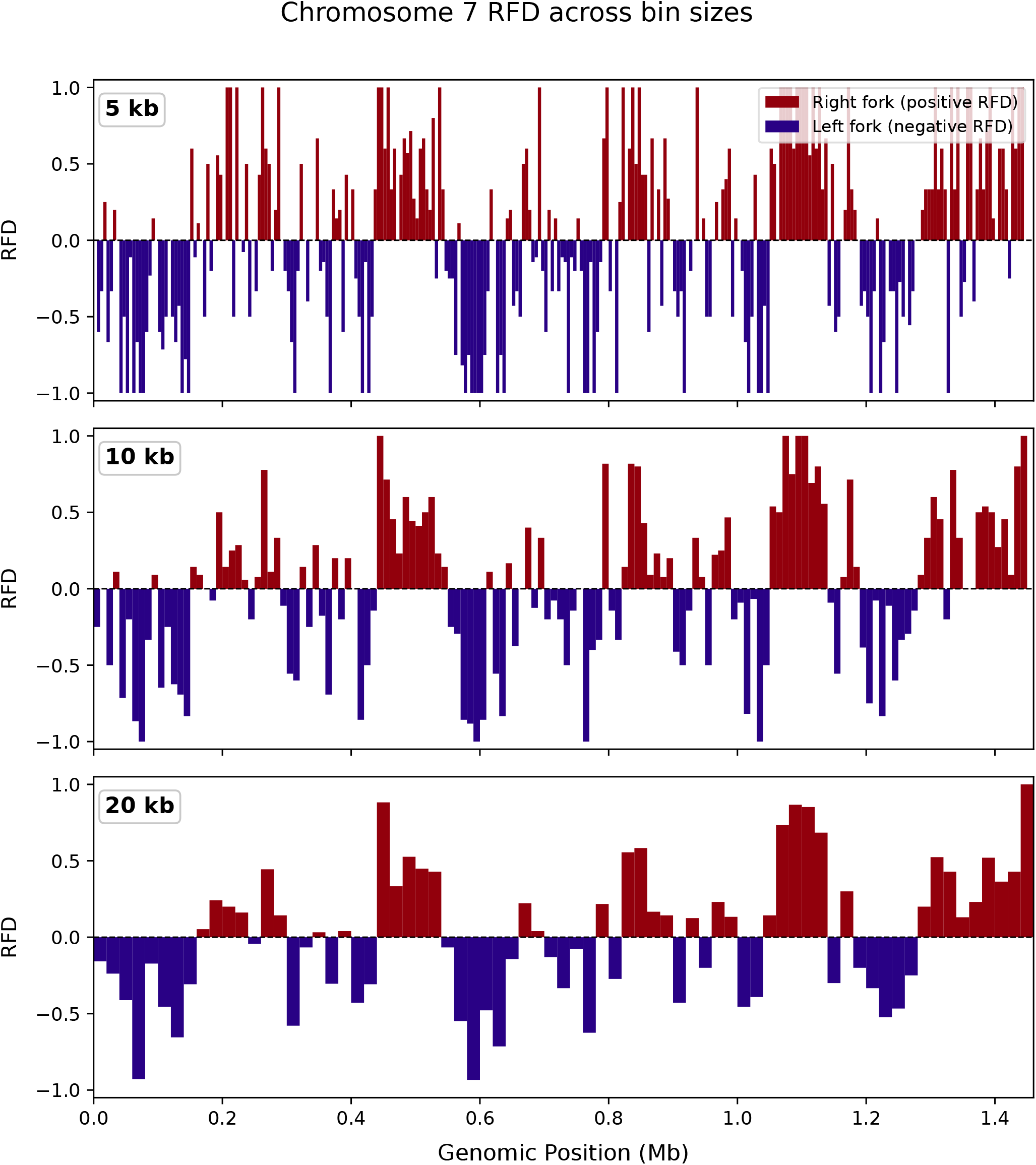
Replication fork directionality for *P. falciparum* chromosome 7 at three different bin sizes: 5 kb (upper), 10 kb (middle), and 20 kb (lower).

**Fig. S2.**
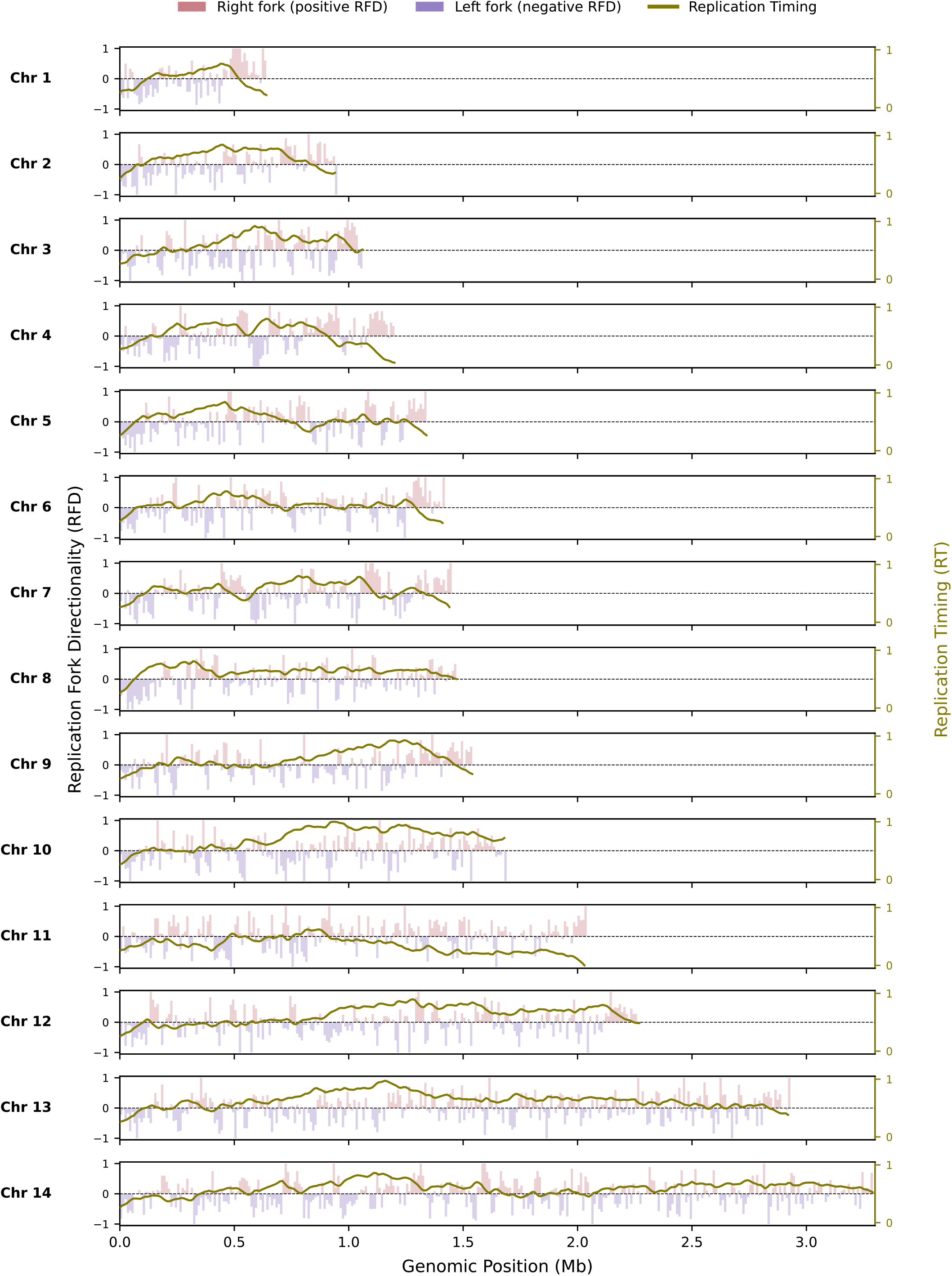
Replication fork directionality (y-axis, left) and replication timing (y-axis, right) for all 14 *P. falciparum* nuclear chromosomes.

**Fig. S3.**
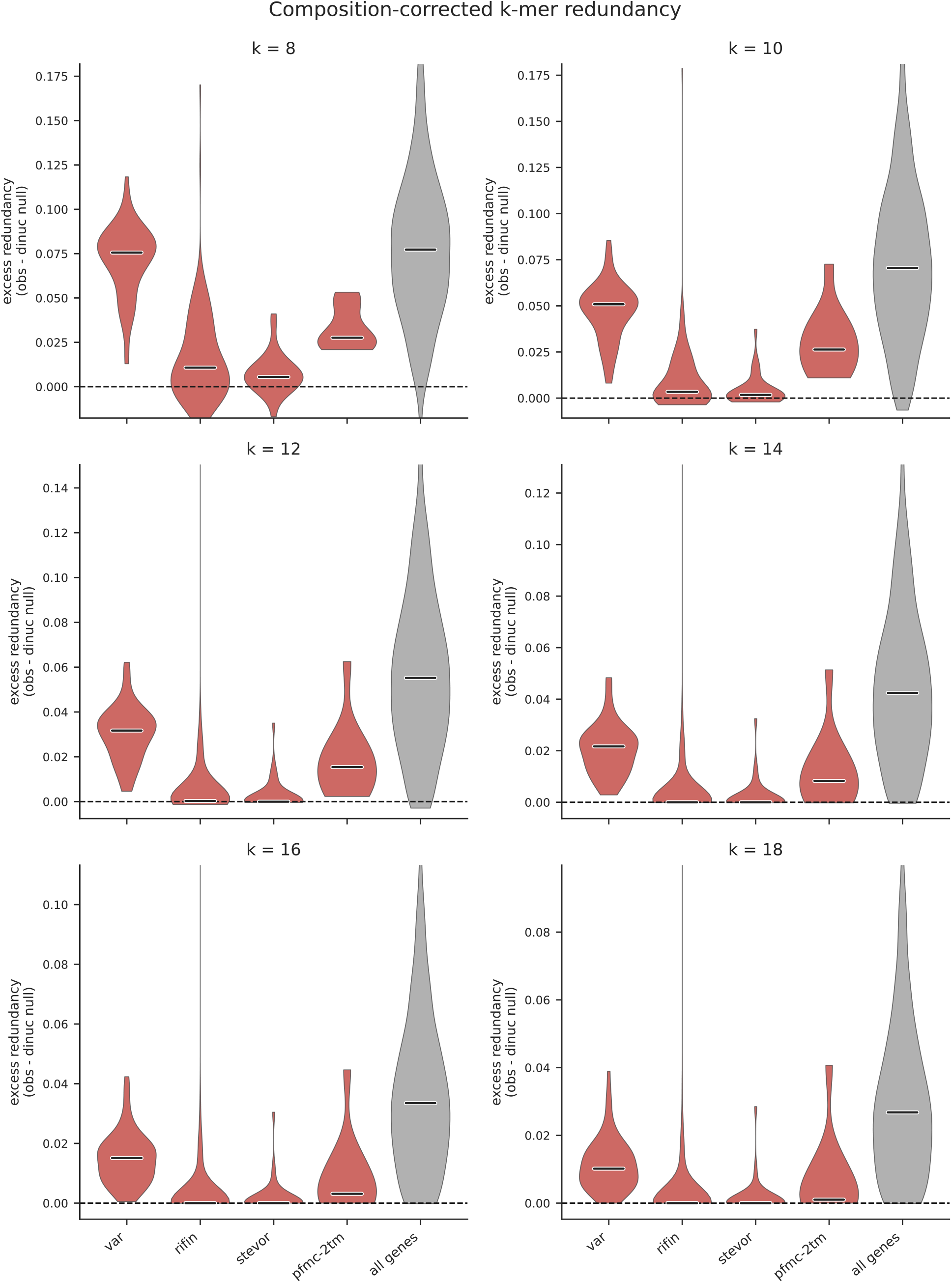
Internal sequence repetitiveness quantified from the *P. falciparum* 3D7 reference genome (assembly ASM276v2). Each protein-coding gene was tiled into non-overlapping 1,000-bp windows. For each window, the *k*mer redundancy *R* = 1 −(distinct *k*mers*/*total number of overlapping *k*mers) was computed so that *R* = 0 indicates a window of all-unique *k*mers and *R →* 1 indicates a highly repetitive sequence. To isolate repeat structure beyond that expected from the AT-richness of each gene family, each gene was compared against a composition-matched null generated by dinucleotide-preserving shuffling with the Altschul–Erikson algorithm, which randomises sequence while preserving mononucleotide and dinucleotide frequencies and the terminal bases. The excess redundancy shown is the redundancy of the gene sequence minus the mean redundancy of the matched null. Results are shown for different values of *k*.

